## Supplementary File 1 for "Altered hierarchical auditory predictive processing after lesions to the orbitofrontal cortex"

**Supplementary File 1a. Mean amplitude centred ± 25 ms around the individual peaks for the MMN and P3a components elicited for the Local Deviance response for the two groups.**

| | MMN – Local Deviance | | | | P3a – Local Deviance | | | |
|---|---|---|---|---|---|---|---|---|
| | Mean amplitude (SD) | | *t* value | *P* value | Mean amplitude (SD) | | *t* value | *P* value |
| | *CTR* | *OFC* | | | *CTR* | *OFC* | | |
| AFz | -7.56 (2.95) | -5.46 (1.40) | -2.467 | 0.021 | 3.65 (1.64) | 1.11 (1.68) | 3.887 | <0.001 |
| Fz | -7.73 (2.96) | -4.84 (1.97) | -2.868 | 0.008 | 3.84 (1.63) | 0.43 (2.51) | 4.166 | <0.001 |
| FCz | -6.80 (3.05) | -5.53 (2.31) | -1.185 | 0.258 | 3.94 (1.87) | 0.73 (1.72) | 4.535 | <0.001 |
| Cz | -6.37 (2.93) | -4.83 (2.31) | -1.475 | 0.153 | 3.34 (2.04) | 0.82 (1.62) | 3.452 | 0.002 |
| CPz | -5.52 (2.48) | -2.97 (2.44) | -2.629 | 0.015 | 2.57 (1.17) | 1.96 (1.32) | 1.255 | 0.222 |
| Pz | -5.63 (2.25) | -1.98 (2.04) | -4.303 | <0.001 | 2.14 (1.37) | 1.13 (1.25) | 1.980 | 0.059 |

Mean amplitude measures in µVolts calculated ± 25 ms around the individual peaks for the MMN (i.e., the most negative peak in a post-stimulus window of 50-150 ms) and P3a (i.e., the most positive peak in a post-stimulus window of 130 -310 ms) components, separately for the healthy control participants (CTR) and the OFC lesion patients (OFC). Standard Deviation (SD) is given in brackets. *P* values as a result of independent samples t-tests comparing the component mean amplitude between the two groups for the midline electrodes.

**Supplementary File 1b. Mean amplitude centred ± 40 ms around the individual peaks for the MMN and P3a components elicited for the Local + Global Deviance response for the two groups.**

| | MMN – Local + Global Deviance | | | | P3a – Local + Global Deviance | | | |
|---|---|---|---|---|---|---|---|---|
| | Mean amplitude (SD) | | *t* value | *P* value | Mean amplitude (SD) | | *t* value | *P* value |
| | *CTR* | *OFC* | | | *CTR* | *OFC* | | |
| AFz | -8.42 (3.19) | -4.78 (3.02) | -2.976 | 0.007 | 7.32 (4.55) | 5.52 (2.95) | 1.169 | 0.254 |
| Fz | -8.71 (4.44) | -6.41 (3.17) | -1.500 | 0.147 | 7.16 (4.03) | 5.08 (3.69) | 1.363 | 0.185 |
| FCz | -8.41 (4.02) | -5.18 (3.98) | -2.054 | 0.051 | 8.05 (4.63) | 7.81 (5.85) | 0.118 | 0.907 |
| Cz | -6.80 (3.60) | -2.28 (3.66) | -3.165 | 0.004 | 7.58 (4.38) | 7.03 (6.81) | 0.247 | 0.807 |
| CPz | -4.99 (3.60) | 0.40 (4.15) | -3.490 | 0.002 | 6.85 (4.18) | 9.11 (5.83) | -1.146 | 0.263 |
| Pz | -5.09 (2.42) | -0.62 (2.73) | -4.432 | <0.001 | 5.37 (4.16) | 6.50 (5.50) | -0.591 | 0.560 |

Mean amplitude measures in µVolts calculated ± 40 ms around the individual peaks for the MMN (i.e., the most negative peak in a post-stimulus window of 50-250 ms) and P3a (i.e., the most positive peak in a post-stimulus window of 150-350 ms) components, separately for the healthy control participants (CTR) and the OFC lesion patients (OFC). Standard Deviation (SD) is given in brackets. *P* values as a result of independent samples t-tests comparing the component mean amplitude between the two groups for the midline electrodes.

**Supplementary File 1c. 50%-area latency for the MMN and P3a components elicited for Local and Local + Global Deviance response for the healthy control participants.**

| | MMN | | | | P3a | | | |
|---|---|---|---|---|---|---|---|---|
| | Latency (SD) | | Diff. (msec) | *P* value | Latency (SD) | | Diff. (msec) | *P* value |
| | *Local* | *Local+Global* | | | *Local* | *Local+Global* | | |
| Fz | 107.31 (07.21) | 127.68 (20.12) | 20.37 | 0.003 | 235.10 (17.69) | 279.88 (21.38) | 44.78 | < 0.001 |
| FCz | 106.61 (07.20) | 127.82 (20.15) | 21.21 | 0.002 | 229.80 (18.19) | 280.02 (26.27) | 50.22 | < 0.001 |
| Cz | 106.06 (06.55) | 128.52 (19.65) | 22.46 | <0.001 | 235.24 (25.29) | 282.95 (28.77) | 47.71 | < 0.001 |
| CPz | 106.47 (06.55) | 130.19 (24.89) | 23.72 | 0.005 | 233.29 (17.45) | 283.79 (24.57) | 50.50 | < 0.001 |
| Pz | 107.31 (05.81) | 132.14 (24.50) | 24.83 | 0.001 | 225.75 (22.77) | 292.58 (33.90) | 66.83 | < 0.001 |

50%-area latency measures in milliseconds (msec) from the onset of the fifth tone for the MMN and P3a components, separately for the Local and Local + Global Deviance responses in healthy control participants. Diff. is the latency difference between the two task conditions (Local + Global vs. Local Deviance) given in msec; *P* values as a result of independent samples t-tests comparing the component's 50%-area latency between the two task conditions. Standard Deviation (SD) is given in brackets.

**Supplementary File 1d. 50%-area latency for the MMN and P3a components for the difference wave (Local + Global minus Local Deviance response) for the two groups.**

| | MMN | | | | P3a | | | |
|---|---|---|---|---|---|---|---|---|
| | Latency (SD) | | Diff. (msec) | *P* value | Latency (SD) | | Diff. (msec) | *P* value |
| | *CTR* | *OFC* | | | *CTR* | *OFC* | | |
| Fz | 164.79 (15.10) | 176.04 (28.36) | 11.25 | 0.210 | 292.07 (28.65) | 346.78 (27.60) | 50.71 | < 0.001 |

| | | | | | | | | |
|---|---|---|---|---|---|---|---|---|
| FCz | 163.96 (15.41) | 184.18 (18.86) | 20.29 | 0.006 | 304.02 (36.42) | 356.38 (50.93) | 52.36 | 0.006 |
| Cz | 163.39 (16.48) | 187.11 (18.33) | 23.72 | 0.002 | 311.41 (33.78) | 349.54 (40.31) | 38.13 | 0.015 |
| CPz | 165.21 (28.56) | 189.15 (17.24) | 23.95 | 0.023 | 302.48 (28.76) | 371.68 (44.08) | 69.19 | < 0.001 |
| Pz | 165.63 (23.99) | 191.83 (21.71) | 26.20 | 0.008 | 306.53 (38.20) | 368.91 (28.09) | 62.38 | < 0.001 |

50%-area latency measures in milliseconds (msec) from the onset of the fifth tone for the difference waves (Local + Global minus Local Deviance response) at the time window of the MMN and P3a components, separately for the healthy control participants (CTR) and the OFC lesion patients (OFC). Diff. is the latency difference between the two groups (OFC vs. CTR) given in msec; *P* values as a result of independent samples t-tests comparing the component's 50%-area latency between the two groups. Standard Deviation (SD) is given in brackets.

### Supplementary File 1e. Characteristics of lesions to the LPFC

| | **Etiology** | **Lesion size (cm$^3$)** | | | **BA (Left hemisphere)** | **BA (Right hemisphere)** |
|---|---|---|---|---|---|---|
| OFC | | Total | L | R | | |
| 1 | Low Grade Glioma | 109.3 | 0 | 109.3 | _ | 4, 6, 8-11, 32, 38, 44-48 |
| 2 | Low Grade Glioma | 0.5 | 0.5 | 0 | 45 | _ |
| 3 | Low Grade Glioma | 158.5 | 0.3 | 158.2 | _ | 6, 8-11, 24, 32, 38, 44-48 |
| 4 | Low Grade Glioma | 28.1 | 0 | 28.1 | _ | 9, 44-46, 48 |
| 5 | Low Grade Glioma | 41.8 | 0 | 41.8 | _ | 9, 10, 32, 44-48 |
| 6 | Frontal Meningioma | 29.9 | 0 | 29.9 | _ | 4-6 |
| 7 | Low Grade Glioma | 33.5 | 33.5 | 0 | 6, 8, 9, 44, 48 | _ |
| 8 | Low Grade Glioma | 12.5 | 12.5 | 0 | 6, 44, 48 | _ |
| 9 | Low Grade Glioma | 51.6 | 0 | 51.6 | _ | 6, 8, 9, 44-46, 48 |
| 10 | Low Grade Glioma | 7.3 | 7.3 | 0 | 9, 10, 32, 46 | _ |

Etiology, size (L, left; and R, right hemisphere), and affected Brodmann Areas (BA) for each hemisphere. The sign "_" is used when no lesion was present in a given hemisphere. Lesions that comprised < 0.2 cm$^3$ in any given BA are not reported.

### Supplementary File 1f. Demographics and neuropsychological performance measures per group.

| **Demographics** | **CTR** | **SD** | **LPFC** | **SD** | ***F* Value** | ***p* Value** | **Stat.** |
|---|---|---|---|---|---|---|---|
| N | 14 | | 10 | | | | |
| Gender (females: males) | 8:6 | | 6:4 | | | | |
| Age years (range) | 47.6 (34-66) | 10.3 | 40.9 (29-65) | 11.8 | 2.29 | 0.15 | ns |
| Education years (range) | 16.1 (13-21) | 2.0 | 15.9 (12-20) | 2.4 | 0.07 | 0.79 | ns |
| **Neuropsychological tests** | | | | | | | |
| Total IQ | 115.4 | 10.3 | 111.8 | 15.1 | 0.48 | 0.50 | ns |
| Digit Span Total | 14.8 | 2.9 | 16.2 | 4.6 | 0.44 | 0.52 | ns |
| Digit Span – Forward | 8.5 | 1.5 | 9.7 | 2.7 | 2.13 | 0.16 | ns |
| Digit Span – Backward | 6.3 | 1.8 | 6.5 | 2.1 | 0.00 | 0.99 | ns |
| Trail Making Test (TMT) | | | | | ***U* Value** | | |
| TMT 2 – Number sequencing | 30.6 | 10.1 | 29.0 | 9.2 | 61.00 | 0.63 | ns |

| | | | | | | | |
|---|---|---|---|---|---|---|---|
| TMT 3 – Letter sequencing | 27.9 | 10.5 | 28.1 | 5.9 | 79.00 | 0.63 | ns |
| TMT 4 – Number-letter switching | 73.8 | 27.1 | 69.3 | 23.8 | 63.50 | 0.71 | ns |
| Color-Word Interference Test (CWIT) | | | | | | | |
| CWIT 1 – Color naming | 31.2 | 6.1 | 30.5 | 78.0 | 61.50 | 0.83 | ns |
| CWIT 2 – Word reading | 22.4 | 3.3 | 22.3 | 5.1 | 61.00 | 0.83 | ns |
| CWIT 3 – Inhibition | 52 | 9 | 55.4 | 17.5 | 75.00 | 0.56 | ns |
| CWIT 4 – Inhibition/switching | 58.2 | 11.9 | 57.6 | 12.4 | 65.50 | 1.00 | ns |
| California Verbal Learning Test (CVLT-II) | | | | | | | |
| Total learning trial 1 – 5 | 57.7 | 12.4 | 57.9 | 9.9 | 63.50 | 0.93 | ns |
| Short-term free recall | 15.2 | 1.2 | 15.5 | 1.0 | 72.00 | 0.69 | ns |
| Long-term free recall | 13.7 | 2.6 | 14.0 | 1.6 | 65.50 | 1.00 | ns |

Comparison of the age, years of education, IQ and Digit Span Test between the two groups (One-Way ANOVA). Comparison of the non-normally distributed raw test scores, Trail Making Test (TMT), Color-Word Interference Test (CWIT) and the California Verbal Learning Test 2nd Edition (CVLT-II) between the two groups (non-parametric independent samples Mann-Whitney U test). Values given are means, with standard deviation (SD). CTR, healthy control group. LPFC, group with lesion to the lateral prefrontal cortex. ns, the statistical test was not significant.

**Supplementary File 1g. Additional measurements that could bias the neural data for the CTR and OFC group.**

| | CTR | SD | OFC | SD | *U* Value | *p* Value | Stat. |
|---|---|---|---|---|---|---|---|
| **Clean trials** | 649.71 | 89.20 | 672.25 | 24.82 | 89.00 | 0.82 | ns |
| Regular block | 334.36 | 56.59 | 333.75 | 24.86 | 75.00 | 0.67 | ns |
| Irregular block | 315.36 | 46.79 | 338.50 | 8.93 | 123.00 | 0.046 | * |
| Control | 280.86 | 51.24 | 281.58 | 20.51 | 75.00 | 0.67 | ns |
| Local Deviance | 267.14 | 38.76 | 285.17 | 8.84 | 113.00 | 0.15 | ns |
| Global Deviance | 48.21 | 9.63 | 53.33 | 1.3 | 115.00 | 0.12 | ns |
| Local + Global Deviance | 53.5 | 5.74 | 52.17 | 4.57 | 76.50 | 0.71 | ns |
| | | | | | ***F* Value** | | |
| **Noisy channels** | 3.00 | 2.11 | 3.83 | 2.92 | 0.71 | 0.41 | ns |
| **Blinks** | | | | | | | ns |
| Regular block | 21.04 | 10.92 | 18.14 | 12.62 | 0.40 | 0.54 | ns |
| Irregular block | 21.58 | 12.94 | 18.18 | 12.84 | 0.45 | 0.51 | ns |
| Control | 18.19 | 10.91 | 19.22 | 13.77 | 0.05 | 0.83 | ns |
| Local Deviance | 22.53 | 14.37 | 21.99 | 16.67 | 0.01 | 0.93 | ns |
| Global Deviance | 20.64 | 12.95 | 14.38 | 9.33 | 1.94 | 0.18 | ns |
| Local + Global Deviance | 23.89 | 12.56 | 17.06 | 11.77 | 2.02 | 0.17 | ns |

Comparison of the number of trials after removing the noisy segments between the two groups (non-parametric independent samples Mann-Whitney U test) and comparison of the number of noisy channels and blinks per minute between the two groups (One-Way ANOVA) across the task blocks (Regular and Irregular) and the experimental conditions (Control, Local Deviance, Global Deviance and Local + Global Deviance). Values given are means, with standard deviation (SD). CTR, healthy control group. OFC, group with lesion to the orbitofrontal cortex. ns, the statistical test was not significant.
